# The midbody and midbody remnant are assembly sites for RNA and active translation

**DOI:** 10.1101/2022.11.01.514698

**Authors:** Sungjin Park, Randall D. Dahn, Elif Kurt, Adrien Presle, Kathryn VanDenHeuvel, Cara Moravec, Ashwini Jambhekar, Olushola Olukoga, Jason Shepherd, Arnaud Echard, Michael Blower, Ahna R. Skop

## Abstract

The midbody (MB) is a transient structure at the spindle midzone that is required for cytokinesis, the terminal stage of cell division. Long ignored as a vestigial remnant of cytokinesis, we now know MBs are released post-abscission as extracellular vesicles called MB remnants (MBRs) and can modulate cell proliferation, fate decisions, tissue polarity, neuronal architecture, and tumorigenic behavior. Here, we demonstrate that the MB matrix—the structurally amorphous MB core of unknown composition—is the site of ribonucleoprotein assembly and is enriched in mRNAs that encode proteins involved in cell fate, oncogenesis, and pluripotency, that we are calling the MB granule. Using a quantitative transcriptomic approach, we identified a population of mRNAs enriched in mitotic MBs and confirmed their presence in signaling MBR vesicles released by abscission. The MB granule is unique in that it is translationally active, contains both small and large ribosomal subunits, and has both membrane-less and membrane-bound states. Both MBs and post-abscission MBRs are sites of spatiotemporally regulated translation, which is initiated when nascent daughter cells re-enter G1 and continues after extracellular release. We demonstrate that the MB is the assembly site of an RNP granule. MKLP1 and ARC are necessary for the localization and translation of RNA in the MB dark zone, whereas ESCRT-III was necessary to maintain translation levels in the MB. Our data suggest a model in which the MB functions as a novel RNA-based organelle with a uniquely complex life cycle. We present a model in which the assembly and transfer of RNP complexes are central to post-mitotic MBR function and suggest the MBR serves as a novel mode of RNA-based intercellular communication with a defined biogenesis that is coupled to abscission, and inherently links cell division status with signaling capacity. To our knowledge, this is the first example of an autonomous extracellular vesicle with active translation activity.

**Highlights:**

- The MB, the center region of the intercellular bridge, is the assembly site of a ribonucleoprotein granule, we call the MB granule
- Distinct oncogenic and pluripotent transcription factor RNAs, including *Jun/Fos* and *KLF4*, are packaged in MBs and MBRs
- The MB granule is coincident with the MB matrix, or dark zone, of the MB
- The Kif23/MKLP1 kinesin is a core hexanediol-sensitive MB granule component
- The MB and MBR are site of active translation that begins in early G1 and continues post-mitotically
- MKLP1 and ARC are necessary for RNA targeting/maintenance and translation at the MB
- Depletion of ESCRT-III increases the levels of translation during abscission
- Abscission releases MBRs as MB granule-harboring, translating extracellular vesicles
- Multiple cell types including cancer, stem, neural stem, all have actively translating MBRs
- MBRs are proposed as a novel mode of intercellular communication by extracellular vesicle-mediated direct transfer of RNA

## Introduction

The midbody (MB) is a protein-rich structure assembled during mitosis at the overlapping plus ends of spindle microtubules, where it recruits and positions the abscission machinery that separates dividing cells(*1–13*). Long thought to be quickly internally degraded in daughter cell lysosomes, recent studies revealed that a majority of MBs are released extracellularly as membrane-bound particles, or extracellular vesicles, following bilateral abscission from nascent daughter cells(*10*, *14–16*). Released post-mitotic MB remnants (MBRs) are bound and tethered by neighboring cells, internalized, and can persist in endosomal compartments for up to 48 hours as signaling organelles (termed MB-containing endosomes or MBsomes) before being degraded by lysosomes(*10*, *12*, *17*, *18*). Distinct cell types, including cancer and stem cells, exhibit differing avidities for internalizing MBRs (*11*, *19*), and exogenous addition of MBRs correlates with increased proliferation and tumorigenic behavior(*11*, *12*, *20*). MBRs have been implicated in specifying apicobasal polarity and lumenogenesis in epithelia(*21*); specifying primary cilium formation(*22*), neurite formation(*23*), and dorsoventral axis formation in *Caenorhabditis elegans* embryos(*24*); and specifying stem cell pluripotency(*25*). The functional importance of MBR signaling in the regulation of cell behavior, architecture, and fate is an emerging field, but the signaling mechanisms are only beginning to be understood.

MB structure and composition suggest mechanistic insights. Proteomic analyses of mitotic MBs and MBRs revealed enriched levels of large numbers (approximately 100) of RNA-binding proteins, ribosomal and translational regulators, and RNA-processing proteins(*1*, *12*, *20*, *26*, *27*), several of which have been implicated in phase-separated condensate formation(*28–32*), but the functional significance was unclear. Given these data, we hypothesized that RNA and ribonucleoprotein (RNP) complexes may play unappreciated structural and/or functional roles in MB biology. Supporting this, a population of polypurine-repeat-containing long non-coding RNAs were localized to the MB(*33*), but the identities and functions of these RNAs remain unknown. In the central core of the MB lies the MB matrix(*4*, *34–37*), a structure of unknown composition. It appears as a prominent electron-dense stripe in electron micrographs(*35*, *38*), similar to other membrane-less organelles(*39–42*); under polarized light it is birefringent(*43*), that is, with a refractive index sharply distinct from the surrounding cytoplasm. Whether RNA plays any role in MB structure or function remains unknown and is the subject of this study.

Here, we further define the structural components, organization, and behavior of MBs throughout their uniquely complex life cycle. Using a quantitative transcriptomic approach, we identified a population of mRNAs enriched in mitotic MBs and confirmed their presence in signaling MBR vesicles released by abscission. We demonstrate that the MB is the assembly site of an RNP granule by analyses of hexanediol sensitivity and fluorescence recovery after photobleaching (FRAP), showing that the matrix exhibits material properties expected of an RNP granule. We show the biochemical activities of MBs are temporally coupled to cell cycle status: MBs initiate translation of stored mRNAs in late telophase as pre-abscission daughter cells re-enter G1 of the cell cycle and continue translation following abscission and MBR release. Last, we found that MKLP1 and ARC play a role in mediating the assembly and maintenance of RNA aggregates and active translation at the MB. In contrast, ESCRT-III is necessary for the modulation of translation levels. We present a model in which the assembly and transfer of RNP complexes are central to post-mitotic MBR function and suggest a novel mode of intercellular communication via extracellular vesicles with defined biogenesis that is coupled to abscission and inherently links cell division status with signaling capacity.

## Results

### Midbodies and midbody remnants are sites of RNA storage

An MB-enriched transcriptome was identified using a comparative genomics approach. Three cell cycle-specific RNA-Seq libraries were prepared from synchronized Chinese Hamster Ovary (CHO) cell populations in interphase, metaphase, and MB/intercellular bridge stage (Fig. 1A). Specifically, whole-cell tubulin, metaphase spindles, and MB spindle microtubules were harvested from CHO cells at interphase, metaphase, and MB stage, respectively, and mRNAs associated with these structures were isolated and sequenced using our previously published methods(*1*). Comparative analysis identified 22 transcripts enriched in the MB stage relative to total mRNAs associated with interphase and metaphase microtubules, with enrichment defined as reads per kilobase million (RPKM) values greater than 1.0 (Fig. 1B; Supp. Tables 1–4). Gene ontology analysis identified that the 22 transcripts encoded factors functioning in transcription, cell fate, cell cycle, RNA processing, and signal transduction (Fig. 1C). A majority of these 22 RNAs are expressed as proteins during late telophase and in the MBR (Fig. S1A). To our surprise, one transcript encoded a critical regulator of cytokinesis, the Centralspindlin component of kinesin Kif23/Mklp1/CHO1(*34*, *44*). Another transcript encoded a member of the TIS11 family of RNA-binding proteins, Zfp36, which has been implicated in regulating RNA stability and RNP granule function in multiple contexts(*45–47*). Perhaps more surprising were the 10 transcription factor-encoding mRNAs identified that are implicated in proliferation, pluripotency, cell fate, cell death, and oncogenesis and that did not have a reported role in cytokinesis (Fig. 1C; Supp. Table 1). In addition, re-evalulation of 99 previously identified RNA binding proteins identified in the midbody proteome (*1*), suggest that these RBPs might perform multiple functions in nucleic acid binding, post-mitotic cell fate functions, cell division, proliferation and development (Fig. S1B-C).

**Figure 1:**
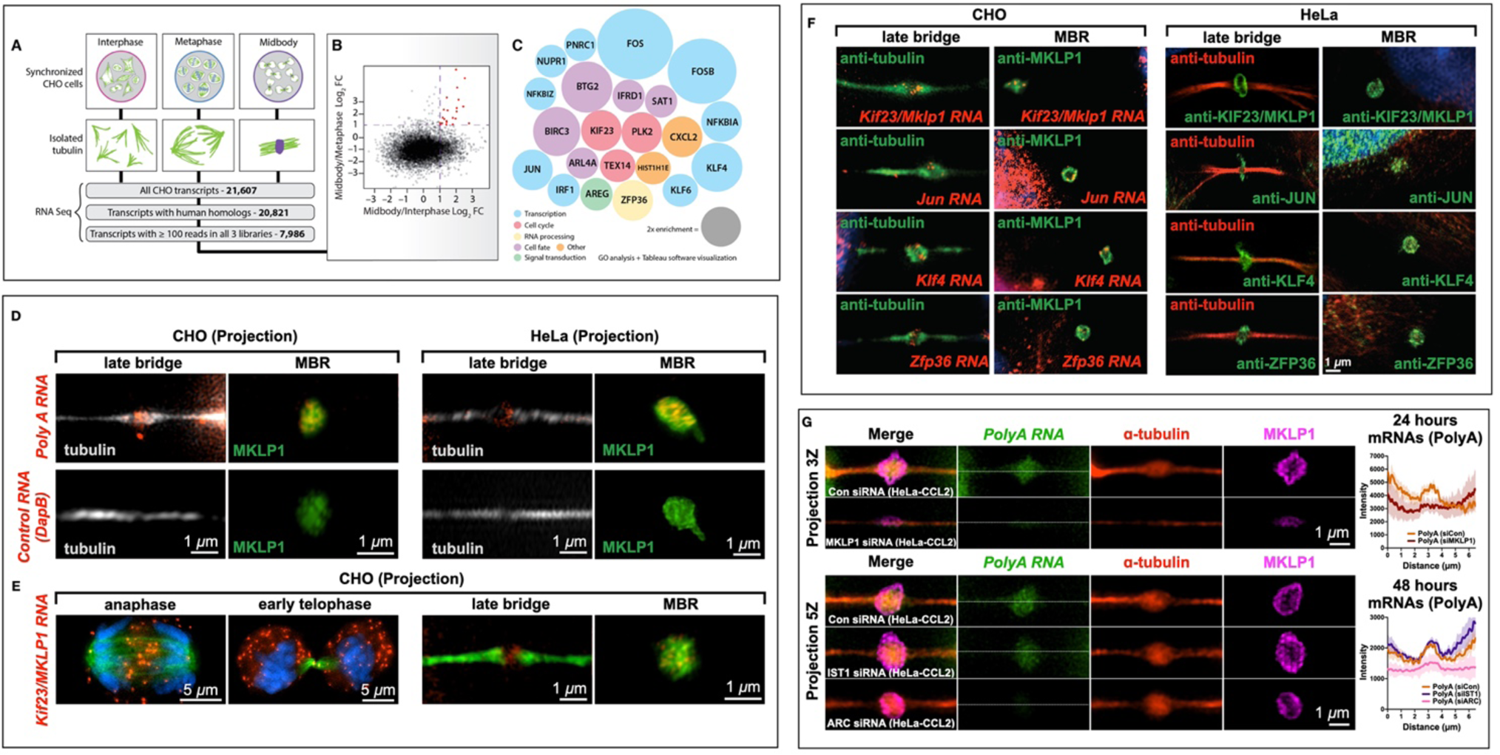
Midbodies and midbody remnants are sites of RNA storage. MKLP1 and ARC are necessary for mRNA localization and maintenance in the dark zone. **(A-C)** RNA-Seq analysis of the MB transcriptome. mRNA was sequenced from three stages of the cell cycle: interphase, metaphase, and late cytokinesis (or “MB stage”). Tubulin structures were purified, and associated RNAs were isolated and analyzed by RNA-Seq. Of 21,607 distinct CHO transcripts identified, 20,821 could be annotated by gene ontology. Of those, 7,986 had ≥100 reads in all cell cycle stages and were further analyzed as plotted in (B). Raw data can be found in Supp. Figs. 2–4. **(B)** Transcripts with ≥100 reads in all three populations were compared and plotted based on their log_2_ enrichment scores (RPKM/RPKM). Dotted lines at x = 1 and y = 1 indicate minimum values for 2-fold enrichment. The 22 transcripts enriched in the MB relative to both interphase and metaphase are highlighted in red. **(C)** Enrichment score (relative diameter) and gene ontology groups of the 22 MB-enriched transcripts; colors correspond to gene ontology biological process terms (Fig. 1; see also Supp. Tables 1–4). **(D)** Single-molecule RNAscope/RNA in situ hybridization revealed mRNA in the MB and released MBRs. PolyA-positive mRNAs (red) localized to mitotic MBs (G1) and post-mitotic MBRs in both CHO and HeLa cells, in contrast with the bacterial *DapB* negative control. **(E)** mRNA encoding Kif23, an MB-resident kinesin required for abscission, localized to the spindle overlap from anaphase through abscission; however, in early telophase, *Kif23/MKLP1* was also found in the cytoplasm in distinct puncta as well as at the MB dark zone. In late telophase (or G1), puncta were found in the dark zone but were also highly enriched in cell bodies; the released MBR contained *Kif23/MKLP1* RNA molecules; tubulin is shown in green. **(F)** mRNAs identified as MB-enriched by RNA-Seq co-localized to the MB and MBR in CHO cells. In HeLa cells, their complementary proteins were localized to the dark zone and the MBR. RNAscope experiments demonstrated that four mRNAs (*Kif23, Jun, Klf4*, and *Zfp36*) localized within the MB matrix, or alpha-tubulin-free zone, of the mitotic MB during G1, and post-mitotically in the MBRs. Proteins encoded by these transcripts similarly localized to mitotic MBs and post-mitotic MBRs in HeLa cells. **(G)** PolyA signals (green) localized to the MB matrix surrounded by *MKLP1* signal (magenta) in HeLa cells. RNAscope fixation techniques led to loss of the MB dark zone as seen by the tubulin bulge along the intercellular canal (red). Quantification of the line scans revealed that loss of *MKLP1* by siRNA knockdown led to a decrease in polyA mRNA in *MKLP1* siRNA-treated cells. Loss of *ESCRT-III/IST1* did not affect RNA levels, but loss of *ARC* led to decreased levels of polyA mRNA in the MB. Scale bars are 1 μm unless noted.

First, we confirmed that the MB is a site of RNA storage by verifying that polyA-containing mRNAs were specifically detected in the MBs and MBRs of CHO cells (Fig. 1D). Similar results were seen in human HeLa cells, suggesting that RNA targeting, and storage are likely to be a general property of MBs (Fig. 1D). Of special interest was the dynamic cell cycle-localization pattern observed for transcripts of *Kif23/Mklp1*, which encode an atypical kinesin motor that is widely used as an MB marker and that critically regulates cytokinesis and abscission(*27*, *48–51*). *Kif23* transcripts were localized to the site of spindle microtubule overlap from early anaphase through late telophase (Fig. 1E), coincident with the localization of KIF23 protein (Fig. S2)(*37*). However, the early telophase pattern was unusual. KIF23 protein is normally found at the MB ring(*48*), but we found that, in early telophase, *Kif23* RNA expression occurred as small puncta found throughout the cell bodies and in two distinct spots adjacent to the dark zone (Fig. 1E, early telophase). Following abscission, *Kif23* transcripts were observed in released MBRs, confirming that these transcripts are present in MBs throughout their life cycle (Fig 1E, MBR).

Next, we selected for further testing three mRNAs from our CHO cell RNA-Seq data that encode distinct classes of proteins (Fig. 1F): the oncogenic transcription factor *Jun*, the pluripotency-regulating transcription factor *Klf4*, and the RNA-binding/RNP granule constituent *Zfp36*(*52–54*). We confirmed that all three mRNAs localized to the MBs and MBRs by RNAscope analysis in CHO cells (Fig 1F, CHO). As with *Kif23*, proteins encoded by each of these mRNAs were also observed in MBs and released MBRs in HeLa cells (Fig. 1F, HeLa).

Last, we identified genes necessary to target or maintain RNA localization to the MB. Using a polyA RNAscope probe in HeLa cells, we found that polyA was enriched in the dark zone of the MB in HeLa (CCL2) cells. PolyA RNA enrichment is dependent on KIF23/MKLP1, a mitotic kinesin(*48*, *51, 55*), and ARC, a repurposed viral-like capsid protein involved in synaptic plasticity and memory(*56*). In contrast, ESCRT-III/IST1, a protein complex necessary for abscission(*57*, *58*) which additionally functions as a RNA-binding protein(*59*, *60*), was not required for the localization of polyA RNA to the dark zone of the MB in HeLa cells(Figure 1G).

These RNAscope data confirmed our transcriptomic findings that MB-enriched mRNA populations were localized to MB structures assembled during mitosis, and that factors necessary for RNA movement were required for their localization to the MB. Combined, our data suggest three novel and testable mechanistic hypotheses: specific mRNAs may be physically sequestered at the MB in RNP granules; MB-targeted mRNAs may be locally translated; these mRNAs and proteins may play an important role in MB function and signaling.

### Midbodies are assembly sites of ribonucleoprotein granules

Several lines of evidence suggest the MB may harbor a phase-separated RNP condensate, given that the MB stores RNA (Fig. 1), is highly enriched in RNA-binding proteins (e.g., staufen, eIF3e, ataxin-2L, PABP, and the 40S and 60S ribosomal proteins)(*1*, *26*, *61*, *62*) is enriched in known RNP granule components, including annexinA11(ANXA11)(*63–65*), and exhibits birefringence(*66*). RNA granules are heterogeneous in composition and function but generally contain solid-like, mobility-restricted structural core components and more labile, liquid-like components that control mRNA flux and translational availability. Yet, it remains unclear how RNA granules are dynamically regulated, assembled, maintained, and disassembled(*67*, *68*).

First, we investigated if midbodies are bona-fide RNA aggregates are reversibly disruptable by challenge with the aliphatic alcohol 1,6-hexanediol, which distinguishes liquid-like assemblies from solid-like assemblies by rapidly dissolving only the former(*69–71*). A 90-second treatment with 7.5% hexanediol was sufficient to disrupt MB matrix in dividing HeLa cells, affecting noticeable but incomplete dispersion of the kinesin KIF23 from its native MB localization (Fig. 2A). When hexanediol challenge was followed by recovery in normal medium in a 0- to 30-minute timed series, KIF23 exhibited progressively wider spatial dispersion, accompanied by reaggregation of increasingly larger assemblies that were usually physically continuous with the native MB (Fig. 2A, T=30). Importantly, the main structural component of MBs—bundled spindle microtubules—was unaffected by hexanediol treatment, suggesting the MB matrix exhibits material properties consistent with a liquid-like assembly, whereas other structural components, such as tubulin, do not. In parallel with our KIF23 results, polyA mRNA also exhibited hexanediol-sensitive dispersion from its normal midzone domain and remained detectable in association with KIF23-positive aggregates, but in complementary domains (Fig. 2B). KIF23 has conventionally been attributed to a structural role in MB-bundling spindle microtubules at the midzone and in assembling abscission machinery at the MB (*48*). However, our data suggest that KIF23/MKLP1 may have an additional role in the positional assembly or tethering of RNA aggregates at the antiparallel microtubule overlap of the spindle midzone, which we observed after short interfering RNA (siRNA) knockdown of MKLP1 (Fig. 1G). To determine if the 1,6-hexanediol sensitive behavior is unique to KIF23/MKLP1, we performed live imaging on GFP-MKLP1 and GFP-MKLP2/KIF20B, a related kinesin-6 family member(*72*). Here, we found that only GFP-MKLP1 was sensitive to 1,6-hexanediol, suggesting that this behavior is unique to this kinesin-6 family member (Fig. 2C). We then used FRAP to determine that KIF23/MKLP1 behaved as a non-mobile component within the native MB granule, as there was very little recovery of MKLP1-GFP fluorescence during the very late stages of cytokinesis (Fig. 2D). This suggests that KIF23 may serve as an immobile kinesin scaffold for the MB RNP granule or MB granule. Our FRAP data were gathered in the context of a native MB within an established RNA granule anchored to microtubules. We interpret these data to suggest that KIF23 behavior is exhibits solid-like behavior in intact, native MBs and liquid-like behaviors when weakly hydrophobic bonds are disrupted with 1,6-hexanediol. This is consistent with a functional role for KIF23 in tethering liquid-like RNP aggregates to the microtubule component of the cytoskeleton.

**Figure 2:**
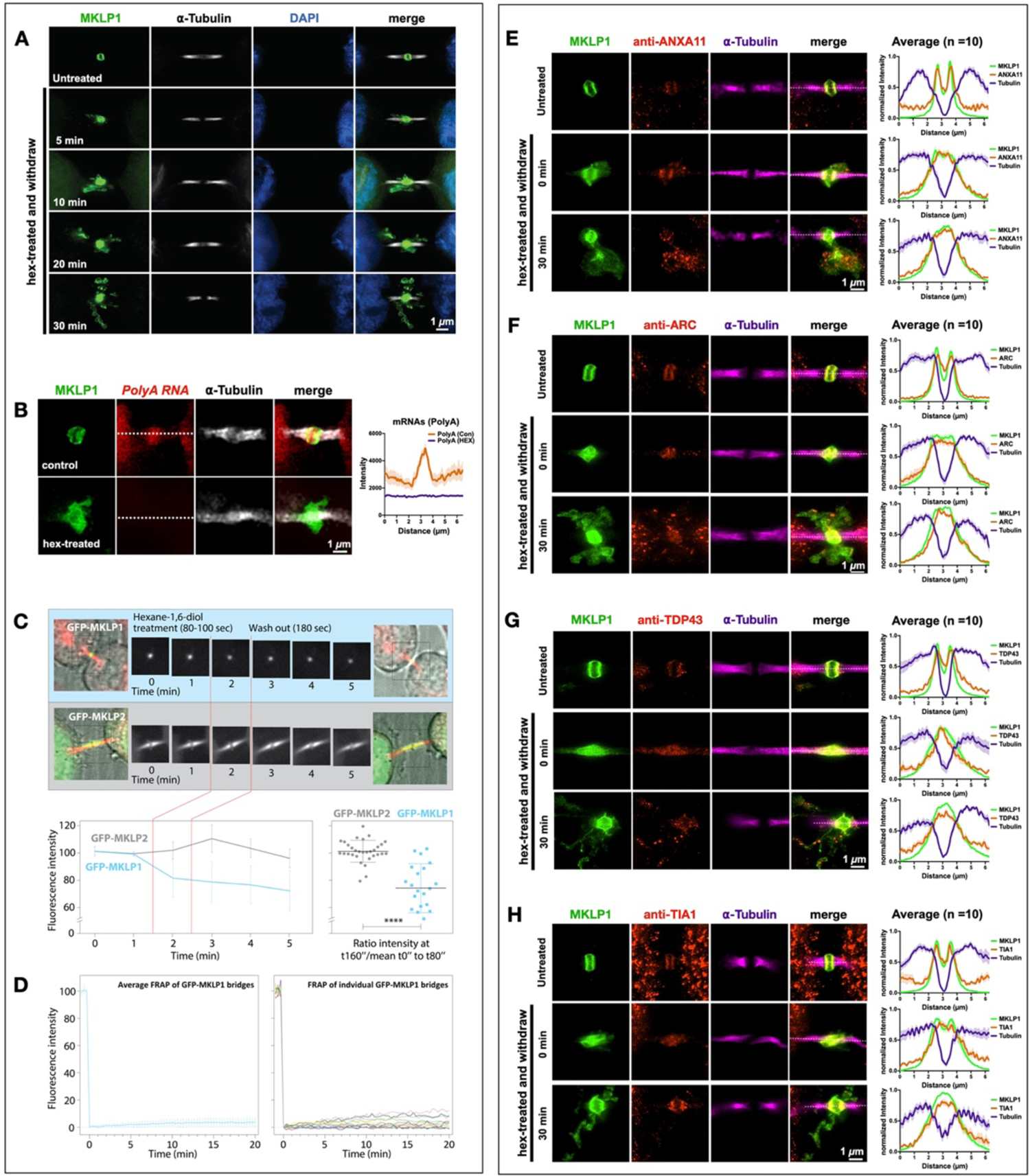
Midbody proteins and RNAs behave as ribonucleoprotein granules. **(A)** Synchronized HeLa cells were treated at the MB stage for 90 seconds with 1,6-hexanediol and then were allowed to recover in normal medium for specified times (T = minutes post-hexanediol). The MB kinesin MKLP1 protein dispersed upon hexanediol addition, reforming spatially disseminated aggregates over time that surrounded the bridge in projected Z-series images. The MB structural component alpha-tubulin was unaffected by hexanediol treatment. **(B)** Treatment with 1,6-hexanediol (hex) also affected polyA localization at the dark zone. We observed a loss of polyA and dissolution of the *MKLP1* signal in the intercellular bridge. **(C)** Live imaging of hexanediol-treated HeLa cells expressing a GFP-MKLP1 fusion protein and incubated with fluorescent SiR-tubulin (red) revealed a rapid and sustained partial loss (30% decrease) of MKLP1 levels at the native MB location; in contrast, the closely related mitotic kinesin MKLP2 fused to GFP exhibited no change in intensity after hexanediol treatment. The 30% loss of MKLP1-GFP after hexanediol treatments reveals that this kinesin is specifically sensitive to 1,6-hexanediol. **(D)** FRAP analysis of GFP-MKLP1 MBs showed no recovery after photobleaching, suggesting little mobility of GFP-MKLP1 within the MB granule in native MBs. **(E-H)**A functional range of MB matrix proteins (ANXA11, ARC, TDP-43, and TIA1) dispersed and reaggregated in apposition to MKLP1 upon hexanediol treatment (T = 0 seconds) and after a long recovery time (T = 30 seconds). Interestingly, all hexanediol-sensitive components tested reaggregated in domains complementary, but tightly apposed, to MKLP1. Of note, we often observed that only a portion of MB factors moves farther away from their original location in the intercellular bridge after hexanediol treatment. For example, the bulk of TIA1 remained diffuse in the dark zone immediately after treatment, but TIA1 quickly assembled back to its normal localization pattern after 30 minutes. MB expression in untreated controls was similar to MKLP1 for all hexanediol-sensitive MB factors (Fig. S2A, B). See also Fig. S4 for a timed series of hexanediol-mediated dissolution and reaggregation of RacGAP, TIA1, ANXA11, and ARC. Scale bars are 1 μm.

Next, we determined whether hexanediol altered the localization of other MB proteins known to function in cytokinesis, as well as putative RNP granule components identified in MBs (Fig. 2E-H; S1C; S2A). In non-treated cells, ANXA11, ARC, TDP-43, and TIA1 all localized to the MB (Fig. 2E-H, controls). After hexanediol treatment and washout, all of the factors tested, were sensitive to hexanediol treatment (Fig. 2E-H, Fig. S2B). Additionally, other MB factors and RNA-binding proteins, including the citron rho-interacting kinase CIT-K, the GTPase RacGAP, and the polyA-binding protein PABP, all of which localized to the MB and MBRs in control cells (Fig. S2A), were hexanediol-sensitive (Fig. S2B). RacGAP, which comprises the Centralspindlin complex with KIF23, formed discrete puncta complementary to KIF23 that resided in KIF23-free pockets directly abutting KIF23 domains (Fig. S2B). Similar patterns of dissolution and reaggregation were observed for two other MB proteins required for cytokinesis, namely CIT-K (Fig. S2B), which directly binds KIF23 and organizes late-stage MB structure, and the phospholipid-binding protein ANXA11 (Fig. 2E), which can tether RNA granules to organellar membranes(*65*). Other midbody factors and RNA-binding proteins also exhibited the same hexanediol-sensitive behaviors (Fig. 2F-H; Fig. S2B). TIA1 localized to the MB matrix and did not appreciably disperse after hexanediol treatment, suggesting it might be an immobile component of the MB structure. Three of these RNA-binding proteins, TIA1, PABP, and TDP-43, function in the assembly and dynamic regulation of stress granules, which are reversible membrane-less organelles that execute cytoprotective defense against environmental stressors by sequestering and translationally silencing mRNAs(*73*, *74*). We also determined that double-stranded RNA, a known extracellular vesicle marker(*75*, *76*), was also located in the MB and MBRs. After hexanediol treatment, double-stranded RNA was found in the cloud of MKLP1 (Fig. S2B, zoomed image). In combination, our data suggest that RNAs targeted to MBs are assembled into phase-separated RNP granules containing mRNA and RNA-binding proteins.

### Midbodies and midbody remnants are sites of localized translation

To determine whether MB mRNAs are translationally activated or silenced, we used two methods to quantify translation. We used the puromycin-based SUnSET technique to label nascent peptides and visualize sites of recent translation using anti-puromycin antibodies(*77*, *78*), and OPP-ClickIT and HPG-ClickIT(*77*), to determine whether active translation occurs in the MB and MBRs. Synchronized HeLa cells pulsed with puromycin for 4 minutes in early telophase (ET) showed little evidence of MB translation(~39% had rings at the ET stage)(Fig.3A, S3A). Parallel cells pulsed just 15 minutes later, in late telophase (LT), showed sharply demarcated toroidal domains of translation encircling the spindle midzone and MB matrix (100% had rings at the LT stage)(Fig.3B; S3A). High levels of translation continued in singly abscised MBs and in doubly abscised and released MBRs (100% had rings)(Fig. 3A, S3A). The puromycin ring during mitosis was also observed in different cell types, including CHO, Retinal Pigmented Epithelial (RPE) cells, Neural Stem/Progenitor cells (NSPCs) (Fig. S3B), We confirmed that this translation signal was indeed active using OPP-ClickIT and HPG-ClickIT(*77*). We observed that the HPG-ClickIT signal gave a hazy disk in the MB in early G1(Fig. 3B). In both singly abscised and doubly abscised MBRs, we saw two distinct regions of HPG-ClickIT signal: a central core and a faint ring of translation around the MBRs that we called the G1 ring (Fig. 3A). Staining with anti-puromycin antibodies revealed a more distinct ring, perhaps owing to the diffusion barrier created by ESCRT (*79*) in the MB, which is located in the same compartment where the bulk of the MB ribosomes and translation regulators are found (Fig. 3C, 40S and 60S)(*36*)

**Figure 3:**
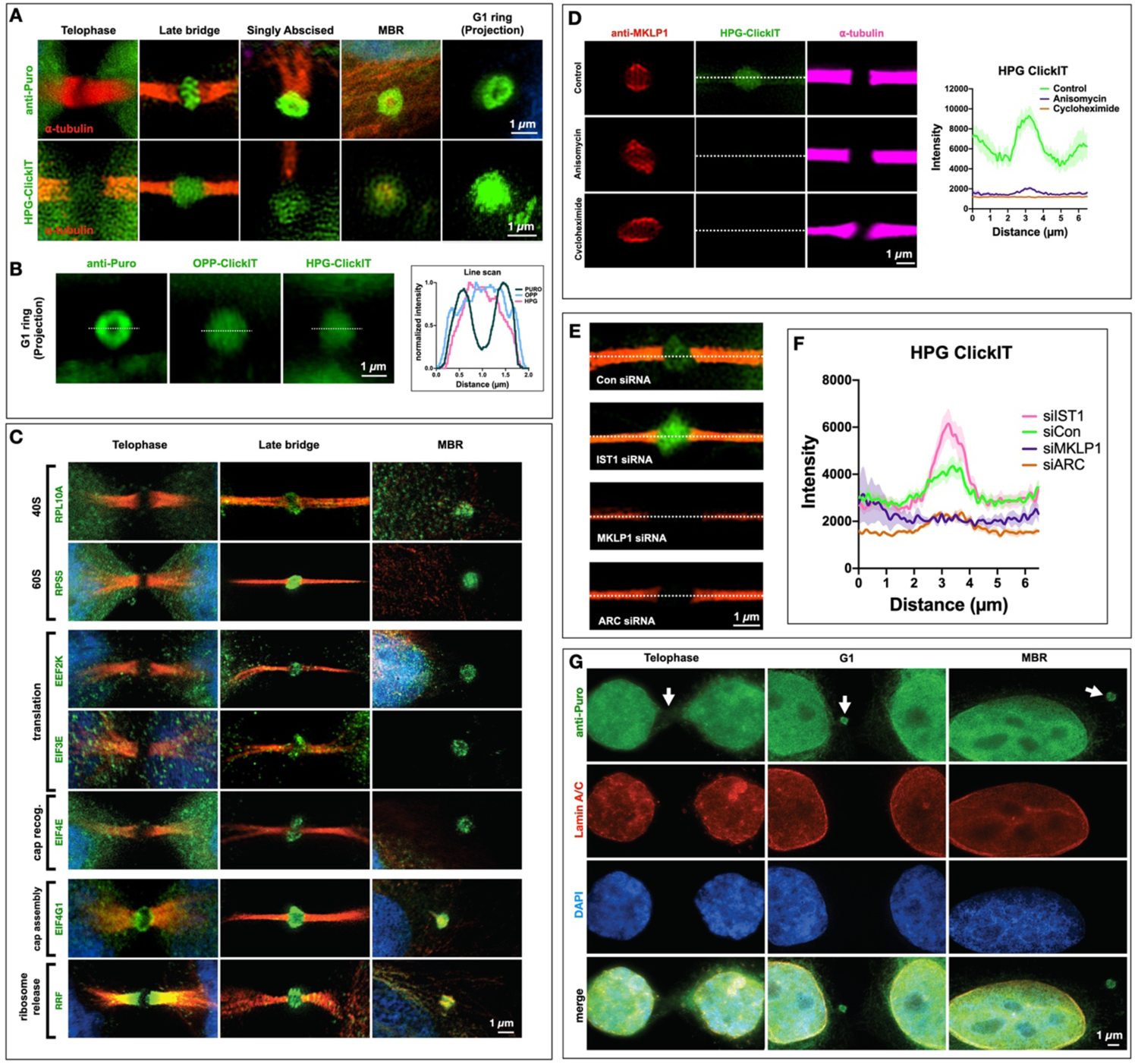
The midbody is a site of spatiotemporally regulated translation. **(A)** SUnSET labeling (α-Puro) revealed that the MB is a translation platform during abscission. Translation was undetected in early MBs (early telophase) but observed at high levels in late MBs (late telophase/G1), in abscising midbodies, and in released extracellular MBRs. Projection revealed that translation occurred in a toroid shape encircling the MB matrix or dark zone. (See Fig. S3A, B for quantification of α-Puro rings per stage). HPG-ClickIT analysis revealed a similar pattern, which suggests that active translation occurred in the dark zone. The HPG-ClickIT pattern appeared as a hazy disk surrounded by a faint ring or cloud. **(B)** The images show the translation patterns from α-Puro (ring) and the OPP-ClickIT and HPG-ClickIT reagents (hazy disk), which indicate a site of recent translation. The graph shows the quantification of the ring and disk patterns. **(C)** Coincident with the puromycin rings, rings were observed for all translation factors previously identified by midbody proteomics (Skop, 2004). Here, 40S and 60S ribosomal subunits (RPL10A and RPS5), translation elongation factors (EEF2K and EIF3E), a cap recognition factor (EIF4E), and a cap assembly regulator (EIF4G1) were first robustly detected in late-stage MBs (abscission/G1) and remained detectable in MBRs. The translational regulators EIF4G1 (cap assembly) and RRF (ribosome release) were present in lateral MB domains in early telophase but re-localized to the translation/ribosome ring at the abscission/G1 transition. **(D)** The robust control HPG-ClickIT signal in the MB dark zone was significantly reduced after treatment with the translation inhibitors anisomycin and cycloheximide. **(E)** Several candidate MB markers were tested, and *ESCRT-III/IST1* was found to regulate the levels of active translation in the MB. *MKLP1* and *ARC* were necessary for translation. Con: control. **(F)** Quantification of the HPG-ClickIT signals in control, *IST1, MKLP1*, and *ARC* siRNA knockdown cells. Con: control. **(G)** Translational onset (α-Puro; arrowheads at MB) occurred precisely as cells formally exited mitosis at the G1 transition, coincident with the mature reformation of the nuclear envelope (detected by lamin A/C) and the de-condensation of chromatin (DNA detected by DAPI staining). DAPI: 4’,6-diamidino-2-phenylindole. Scale bars are 1 μm unless noted.

We treated MBs with anisomycin or cycloheximide to inhibit translation(*77*). We observed that the HPG-ClickIT signal was abolished after these drug treatments and the MKLP1 localization was often distorted (Fig. 3D), suggesting that active translation during late telophase might be necessary for the proper maintenance of MB structure.

### MKLP1, ARC, and ESCRT-III regulate translation in midbodies

To determine which genes might be necessary for the unique translation event that occurs in MBs, we knocked down *ESCRT-III/IST1*, *MKLP1*, and *ARC* using siRNAs in the HeLa (CCL2) cell line (Fig. 3E, S4A). Surprisingly, suppression of ESCRT-III/IST1 led to a sharp increase in active translation in the MB (Fig. 3E-F, S4A). However, in MKLP1 and ARC siRNA-treated cells translation was entirely abolished at the MB (Fig. 3E-F, S4A), suggesting either that these genes are required for translation or that they are required to target or maintain MB RNA. In both parental HeLa (CCL2) and MKLP1-GFP HeLa expressing cell lines, we only observed failure of cytokinesis (bi-nucleate daughter cells) in MKLP1 siRNA-treated cells (Fig. S4B), whereas ESCRT-III/IST1 led to delays in abscission (% MB bridge, Fig. S4C). We favor the latter suggestion, as loss of MKLP1 and ARC led to a loss of RNA signal in the MB (Fig. 1G). Additionally, of the siRNAs we knocked down, only the loss of MKLP1 led to a thinning of the microtubules in the midbody (Fig. S4A), suggesting that in MKLP1 siRNA-treated cells there may be a limited ability to target RNAs.

We also made an unexpected finding that MKLP1 may mediate global translation events, as HPG-ClickIT levels were increased at the MB dark zone in the MKLP1-GFP cell line when compared with the HeLa (CCL2) cell line (Fig. S5A-B).We further observed significant translation throughout the cell bodies and the MB dark zone in the MKLP1-GFP cell line (Fig. S5C). These data suggest that MKLP1 may mediate translation in distinct cellular sites (in the cell body and the MB), and these data represent a caution to others that use of this MKLP1-GFP cell line could confound their results. We also confirmed using HPG-ClickIT that active translation levels changed in the MKLP1-GFP cell line(*80*). We found that knockdown of *ESCRT-III, MKLP1*, and *ARC* by siRNAs had similar effects on translation in the MKLP1-GFP cell line. Loss of ESCRT-III led to increased levels of translation, and MKLP1 and ARC appeared to be required for translation (Fig. S4A). Overall, our finding that localized translation in the MB initiated prior to daughter cell separation raises the possibility that assembly of the MB RNA granule and translation of its contents may be a necessary step during the late steps of cytokinesis or abscission. In addition, this is the first example, to our knowledge, of an autonomous extracellular vesicle with active translation activity, and may reflect a transition in the life stage of the MB RNA granule that is critical to post-mitotic MBR signaling function.

A primary function of many RNP membrane-less compartments such as stress granules is to regulate the translational availability of mRNAs by reversible partitioning into translationally silenced condensates(*39*, *81–83*). Although it is accepted that global translation is severely restricted during mitosis, MB and MBR RNA interference screening and proteome analysis suggest the presence of large complements of both 40S and 60S ribosomal subunit proteins(*1*, *12*, *26*) and translation initiation and elongation factors(*1*, *12*, *26*, *84–86*); we confirmed MB and MBR localization of these proteins in representative samples (Fig. 3C).

### Translation starts at the M/G1 transition

Dividing daughter cells formally exit mitosis, while still joined by the intercellular bridge containing the MB and undergo abscission only after re-entering the G1 phase of the cell cycle when they resume global protein synthesis(*87–89*). We used SUnSET staining(*78*) to determine the relative timing of MB translation initiation with three hallmarks of the M/G1 transition: re-initiation of global translation, nuclear envelope reassembly, and chromatin decondensation. In late telophase, newly segregated chromosomes are fully condensed, the nuclear envelope is beginning to reform, and translation in the MB and daughter cell body was almost undetectable (Fig. 3G, telophase). As daughter cells progress into G1, chromatin decondensation initiates as the nuclear envelope becomes continuous, and active MB translation was observable in the intercellular bridge (Fig. 3G, G1 phase). Following abscission, the euchromatin of interphase daughter cells was observable within fully formed nuclear envelopes, and actively translating extracellular MBRs were visible on plasma membrane surfaces (Fig. 3G, MBR). We, therefore, hypothesize that the G1 transition triggers a burst of translation in a juxta-granular compartment of the MB.

Supporting this hypothesis, we found that most proteins encoded by MB-enriched mRNAs (identified in Fig. 1C) were first detectable in the MB only after the G1 transition (n=10/12; Fig. S1A, C) In telophase, the two cytokinesis factors, KIF23 and TEX14 are seen in the MB matrix and flanking arms of the intercellular bridge, respectively. In contrast, the remaining 10 proteins, which have no reported role in cytokinesis, were undetectable in early telophase, except KLF4, despite being readily seen in late telophase (Fig. S1A); these factors included five transcription factors (JUN, cFOS, FOSB, KLF6, and IRF1), the transcriptional inhibitor IKBalpha, the RNA granule component ZFP36, histone HISTH1, and the multifunctional BIRC3 protein. In contrast, all 12 factors were readily detected in the MB at later stages following transition into G1 and remained detectable in post-abscission MBRs. These data strongly suggest that mRNAs targeted to the MB RNA granule become translationally available coincident with the M/G1 transition and may reflect a critical life cycle transition as the mitotic MB matures toward release as an extracellular MBR with post-mitotic signaling functions.

### ARC leads to a decrease in RNA and translation at the MB

To identify which MB RBP might be responsible for the assembly or maintenance of RNA and the translation activity in the MB, we took a close look at our previously published MB proteome (Fig. S1B). Here, we identify several candidates that might be important for this function, which include TIS11B, Staufen/Stau, Annexin aII, Ataxin 2L and ARC, all of which localized to the MB during G1/late telophase of the cell cycle (Fig. S1C). Using siRNA knockdown, we observed that the PolyA signal at the MB was found in all of our knockdowns except ARC (Fig 1G; Fig. S6A). There was a slight decrease in PolyA signal in TIS11b siRNA-treated cells (Fig. S6A). However, ARC was the only factor necessary to a decrease the HPG-ClickIT signal (Fig. S6B), suggesting that ARC is critical for RNA maintenance and translational activity in the MB.

## DISCUSSION

Until recently, the MB was thought to regulate assembly of the abscission machinery during cytokinesis and then be immediately degraded following cell separation. However, studies in the last decade have demonstrated that post-mitotic MBRs are released by abscission as extracellular vesicles, are internalized to form signaling MBsomes in target cells, and may contribute to driving highly proliferative fates such as tumor and stem cells(*6*, *11*, *12*, *19*, *26*, *34*, *90–92*). In support of this idea, MBRs are preferentially accumulated in tumor and stem cells, and exogenous MBRs can upregulate proliferation-promoting genes, the proliferative index, and anchorage-independent invasiveness(*12*, *20*, *93*). Although the functional importance of post-mitotic MB signaling has been established, the underlying mechanisms remain poorly understood. Recent advances identify a requirement for integrins and EGFR receptor tyrosine kinase signaling in MBsome function; however, this simple model does not sufficiently account for the strikingly large size of MB derivatives nor their structural complexity and multi-stage life cycles. In this study, we characterized the structural components of MB derivatives to gain insight into post-mitotic MBR signaling mechanisms. Importantly, we demonstrated that the MB is the assembly site of an RNP granule that is packaged and released within a large 1- to 2-μm extracellular vesicle following the terminal stages of cell division. We used a transcriptomic approach to characterize specific mRNA populations that are enriched at the MB in a translationally quiescent granule called the MB granule and demonstrated that local translation of MB granule mRNAs was initiated as cells exit mitosis and MBRs are released by abscission. By identifying ongoing translation in MBRs, we provide the first demonstration of active translation in an extracellular vesicle, which implies that dynamic translational availability of MB granule mRNAs may play an active role in subsequent target-cell binding and/or signaling by MBRs.

The reversible formation of RNA granules is the primary mechanism by which cells control translational availability and localization of RNAs to rapidly respond to changing cellular demands. During mitosis, RNA and ribosomal protein sequestration in condensates facilitates global shutdown of protein synthesis and regulates cytoplasmic partitioning. This study identified a novel subtype of mitotic RNA granule with localized assembly at the overlapping spindle microtubules that define MB positioning; thus, we called it the MB granule. The locations of the MB granule and MB matrix precisely correlate. We report that MB granules are also exquisitely sensitive to hexanediol in vitro, a behavior typical of liquid-like assemblies; MB granule-associated RNA-binding proteins and RNA dispersed almost immediately upon hexanediol treatment and then progressively reaggregated in heterotopic puncta and broad clouds continuous with the native MB after hexanediol removal. Interestingly, RNA and RNA-binding proteins reaggregated in domains complementary to reaggregating KIF23, revealing organization within the reforming liquid-like assembly. KIF23 has previously only been suggested to function in microtubule bundling and vesicular trafficking to the MB(*48–50*, *90*, *94*, *95*), so its liquid-like behavior was not predicted, especially as spindle microtubules were unaffected by hexanediol treatment. Somewhat paradoxically, FRAP analysis indicated that MKLP1-GFP was an immotile component of MB granules. We interpret the data to suggest that KIF23 performs a tethering function in MB granules by binding microtubules with its N-terminal motor domains and by binding RNA or RNA-binding protein assemblies with the predicted intrinsically disordered regions near its C-terminus. It remains possible that the heterotopic material observed reflects formation of hexanediol-induced stress granules(*69*) or other aberrant granules caused by prolonged hexanediol exposure(*96*). We think this scenario is unlikely, as we could never detect the stress granule marker G3BP with any of several antibodies tested (data not shown), and our 90-second hexanediol treatments were far below the 50-minute threshold reported for cytotoxic granule induction(*96*).

A basic function of RNA granules is to translationally silence phase-separated mRNAs, and MB granule transcripts are subject to cell cycle-entrained translational control. Proteomic and immunofluorescence analyses revealed that both mitotic MBs and post-mitotic MBRs harbored large quantities of ribosomal proteins and translational regulators. As translation is largely silenced during mitosis, it is perhaps expected that no translation was detectable in telophase-stage MBs or intercellular bridges. Concomitant with the nascent daughter cells re-entering G1 of the cell cycle and reinitiating global translation, a hazy disk and ring of translation was observed within the MB granule but not appreciably in flanking regions. Translational onset temporally prefigures abscission, so it is tempting to speculate that local translation is required for terminal cell separation; however, we have been unable to generate evidence supporting this hypothesis. Active translation persists as post-abscission MBRs are released as extracellular vesicles, strongly implying a post-abscission function for ongoing protein synthesis. MBRs have been shown to enrich phosphatidylserine in their outer membrane leaflets only subsequent to abscission, as they mature toward their ultimate fate of engulfment and MBsome signaling. We suggest that active translation may play a parallel role in maturation of MBRs that potentiates recognition and engulfment by target cells and/or mediates MBsome signaling(*10*, *12*, *20*).

A central problem in the field of MB biology is that the molecular mechanisms underlying MBR and MBsome signaling remain poorly understood. One possible impediment is that MBRs had simply not been conceptualized as extracellular vesicles until more recently(*20*) and did not benefit from the intense research interest focused on extracellular vesicle-mediated intercellular communication, including by direct RNA transfer. In this work, we newly identify MBRs as a unique subtype of extracellular vesicles with several distinguishing features: stochastic biogenesis in mitosis that inherently links cell division status with intercellular signaling; a complex life cycle with both membrane-less and membrane-bound stages; a cargo comprised of a selectively loaded RNA granule; active translation of mRNA cargo; and an extremely large carrying capacity that is greater than 1000-fold more than exosomes on average. We showed that mRNAs were assembled into an MB granule in association with KIF23 and were lost following hexanediol-induced disruption. Our data strongly support the hypothesis that MBRs and internalized MBsomes can signal, at least in part, by the direct transfer of MB granule components. It is also possible that MBR-mediated transfer of RNP complexes is specific to HeLa cells. However, we observed that HeLa cells, CHO, retinal pigment epithelium cells, and neural stem cells harbored MKLP1-positive MBs with puromycin-labeled rings that appeared in G1 (Fig. S3B). We favor the hypothesis that MB granule-mediated RNA transfer is a signaling mechanism fundamental to all cells that divide using an MB, and that MBRs are selectively loaded with distinct transcriptomes in a cell type-specific manner and use ARC, a viral-like capsid(*56*), to facilitate the mechanism of cell-cell communication in all cell types, not just neurons. These hypotheses are readily testable.

We propose a model of the MB life cycle that frames its complex structural dynamism in terms of a novel post-mitotic signaling function: intercellular communication via extracellular vesicle-mediated transfer of RNA (Fig. 4, model). During anaphase of mitosis, selected mRNAs and MKLP1 coacervate and are tethered to overlapping regions of the antiparallel spindle microtubules by a process involving KIF23/MKLP1. As spindle microtubules constrict into an intercellular bridge during early telophase, individual coacervates coalesce into a single large RNP granule at the midzone that we called the MB granule. As nascent daughter cells transition to G1, peri-granular translation initiates throughout the MB and outward toward the ribosome-rich ring, presaging the bilateral assembly of the abscission machinery. Following scission and MBR release as a membrane-bound extracellular vesicle, severed microtubules depolymerize and domains of translation are radicalized around the MB granule as the maturing MBR surveys putative target cells for docking sites. Bound and internalized MBRs evade degradation and persist as MBsomes, releasing MB granule constituents into the recipient cell’s cytoplasm. We suggest that liberated MB granule RNAs are a critical functional component of MBsome signaling in recipient cells that act as templates for direct translation of effector proteins or as templates for epigenetic silencing or as a combination of these two mechanisms.

**Figure 4.**
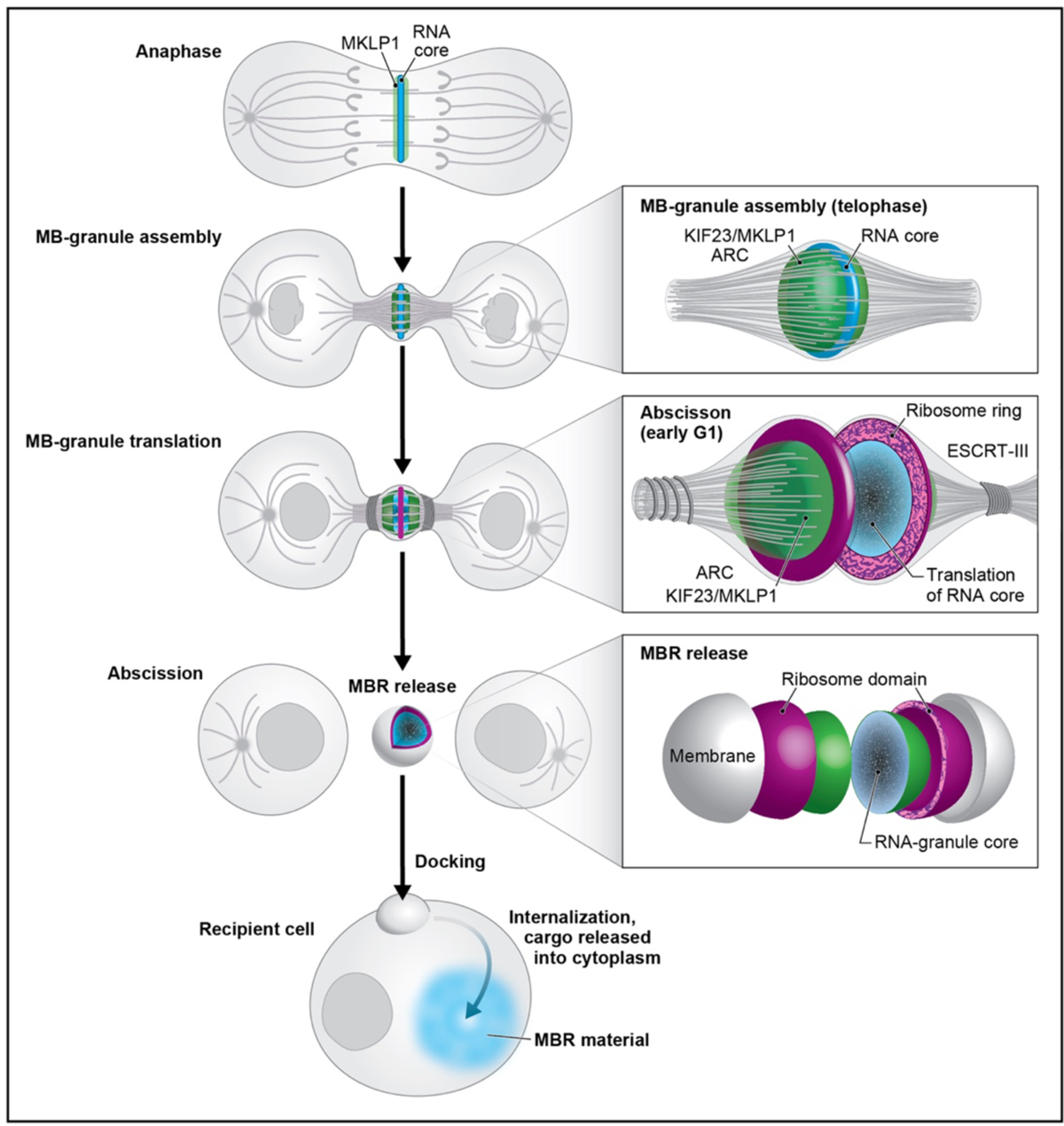
Model of the unique life cycle of the midbody granule and biogenesis of the midbody remnant, a unique actively translating extracellular vesicle. We present a model in which the MB not only plays its traditionally considered role in abscission but also mediates a novel form of intercellular communication. In anaphase, MB-targeted RNAs and associated RNA-binding proteins, such as MKLP1/KIF23 and ARC (both green), begin to form small phase-separated RNP condensates (blue) at the spindle microtubule overlap. Actomyosin ring constriction drives intercellular bridge formation and accretion of a single large MB granule in telophase. At the abscission/G1 transition, ribosomes (magenta) and translation factors surround the RNA core (blue). Translation is active throughout the entire MB granule (blue) and is followed by assembly of the abscission machinery and scission. The MBR is released, which harbors an MB granule core surrounded by a shell of active translation. We propose that MBRs dock to and are internalized by recipient cells, and this process is followed by the transfer of MB granule cargo, including RNA, across endolysosomal membranes into the cytoplasm. We hypothesize that the instructional information resides in the MB granule RNA and serves as templates for either direct translation or epigenetic modulation.

## Author Contributions

SP, RD, AJ, AE, MB, and ARS conceived and designed experiments. RD, SP, AP, KvD, AJ, MB, and AP performed experiments, bioinformatic analyses, data analyses, and data visualization. SP, RD, and ARS co-wrote the manuscript. EK assisted with imaging and figure preparation. CM performed imaging data analysis for Figure 2 with RD. AE and MB contributed to text and figure revisions. JS contributed to info about ARC and antibody validation.

## Acknowledgments

The authors thank Judith Kimble, Robert Singer, Michael Sussman, Harmit Malik, Diana Chu, Abby Dernburg, Karen Schindler, Daniel Jung, Maureen Barr, Dana Miller, and Premal Shah, for sharing their expertise, support, and advice. We are indebted to the work performed by Jessica Shivas, Jennifer Gilbert, and Lan Qin during the initial stages of this project. Special thanks to Chris Morrow and Darcie Moore for the NSPC cells; Sakae Ikeda and Akihiro Ikeda for the RPE cells. Special thanks to Kurt Weiss, Elle Grevstad and Peter Favreau for their technical assistance with structured illumination microscopy at the University of Wisconsin–Madison Biochemistry Optical Core. We can’t thank Adam Steinberg enough (artforscience.com) for pushing the science further by showing gaps in our knowledge throughout the process of our research in our model.

## Materials and Methods

### Cell culture maintenance and synchronization

Chinese Hamster Ovary (CHO) cells (ATCC^®^ CCl-61^™^) were maintained at 37°C and 5% CO2 in DMEM/F-12 (Thermo Fisher, Cat# 11330057) with 10% FBS (Thermo Fisher, Cat# 26140079) and 1% Penicillin-Streptomycin (Thermo Fisher, Cat# 15140-122). “Interphase” CHO cells were asynchronous populations cultured for 48 hours before RNA isolation. Synchronized CHO cell populations were grown as described (Skop, 2004)(*1*): cells were blocked in S phase by two rounds of growth for 16 hours in medium supplemented with 2 mM thymidine (Sigma, Cat# T1895-5G) interrupted with 8 hours incubation with DMEM/F-12 medium. Following the second thymidine block, cells were released into DMEM/F-12 medium for 5 hours and then treated with 100 ng/ml nocodazole (Sigma, Cat# M1404) in DMEM/F-12 medium for 4 hours to arrest cells in metaphase. Mitotic cells were isolated by mechanical shake-off and transferred to DMEM/F-12 medium. “Metaphase” samples were incubated for 15 minutes to allow mitotic spindles to reform, and spindle-associated RNAs were isolated. Following nocodazole wash-out, “MB” samples were incubated for 30–45 minutes until contractile rings were apparent (late telophase/G1), and MB-associated RNAs were harvested. To harvest stage-specific RNAs, interphase microtubules, metaphase spindles, and MBs were isolated as described (Skop, 2004)(*1*). HeLa cells (CCL-2; ATCC) were cultured at 37°C and 5% CO2 in DMEM/high-glucose/GlutaMAX medium (10564029; Thermo Fisher Scientific) supplemented with 10% fetal bovine serum Thermo Fisher, Cat# 26140079 and 1% penicillin-streptomycin (Thermo Fisher, Cat# 15140-122). HeLa cells were synchronized using a similar double thymidine-block procedure, as previously described (*97*). HeLa cells were synchronized to arrest in prophase by culture in 50 ng/ml nocodazole in DMEM/high-glucose/GlutaMAX medium for 16 hours. The mitotic cells were harvested by shake-off, centrifugation (200g/1000rpm by Eppendorf centrifuge 5702, 1min), and release from high-precision cover glasses (Zeiss, Germany, Cat# REF# 0109030091) with pre-warmed DMEM/high-glucose/GlutaMAX medium (90 min, early midbody during telophase; 4 hours, late midbody during G1). The cells were treated with 91 μM puromycin (Sigma-Aldrich, Cat# P8833) in DMEM/high-glucose/GlutaMAX medium for 4 minutes before fixation. Primary hippocampal mouse neural stem cells (NSCs) were isolated by extracting and dissociating hippocampi from 3-5 mice roughly 6 weeks of age, as described previously (Moore et al., 2015). GFP-MKLP1 and GFP MKLP2 HeLa cells (*98*), were cultured at 37°C and 5% CO2 in DMEM/Glutamax (#31966; Gibco, Invitrogen Life Technologies) supplemented with 10% FCS, 1% penicillin-Streptomycin and kept under G418 (40 μg/mL, Gibco). NSCs were cultured at 37°C/5% CO2 in serum-free media: DMEM/F12 GlutaMax (10565018; Invitrogen) with B27 (1:50, 17504044; Invitrogen), penicillin-streptomycin-fungi-zone (1:100, 15140122; Invitrogen), 20 ng/mL FGF-2 (100-18B; PeproTech), EGF (AF-100-15; PeproTech) and 5ug/mL Heparin (H3149; Sigma), as previously described (Morrow et al., 2020). RPE-1 (ATCC^®^ CRL-4000^™^) was cultured in DMEM/F12 (Thermo Fisher) supplemented with 10% fetal bovine serum and penicillin/streptomycin at 37°C in an atmosphere of 5% CO_2_.

### CHO midbody RNA purification and Illumina library preparation

CHO microtubule pellets (interphase, metaphase, and MB stage) were resuspended in approximately 100 μl phosphate-buffered saline (PBS). RNA was purified from each sample using a Qiagen RNeasy kit. PolyA RNA was purified from 1 μg RNA from each sample using an Exiqon LNA dT purification kit in accordance with the manufacturer’s instructions. PolyA RNA at the final purification step was eluted using Illumina Elute/Prime/Fragment buffer. Illumina RNA libraries were constructed using the Illumina TruSeq RNA Sample Preparation Kit v2 in accordance with the manufacturer’s instructions. Each library was barcoded and sequenced on an Illumina HiSeq 2500 system.

### Data deposition

RNA sequences associated with this study have been deposited into the National Institutes of Health Sequence Read Archive (Bioprojects: PRJNA191571 and PRJNA247381).

### Annotation assignment and RNA-Seq data filtering

RNA-Seq reads were collapsed into unique reads using a custom Perl script(*99*). Unique reads were aligned to the RefSeq sequences for the Chinese hamster (*Cricetulus griseus*) using Bowtie 2(*100*). Reads mapping to transcripts were quantified using a custom Perl script(*99*) or HTSeq(*101*). For comparison analyses, we only considered genes with at least 100 reads in all three libraries. After alignment, hamster orthologs were identified using BLASTx (National Center for Biotechnology Information); annotations were automatically assigned using DAVID (https://david.ncifcrf.gov/) and PANTHER (www.pantherdb.org) and then manually curated using gene ontology terms and the UCSC Genome Browser database. Enrichment scores were defined as the ratio of normalized read counts (in RPKM) between libraries; all comparative quantitative analyses of RNA levels were performed using RPKM values or reads per million values. RNA-Seq resulted in 21,607 transcripts, 20,821 of which had human orthologs, with at least one read in any of the three libraries (interphase, metaphase, and MB), resulting in 15,636 transcripts in the interphase library, 17,813 transcripts in the metaphase library, and 16,528 transcripts in the MB library. After low-abundance reads were discarded, 10,424 entries remained in the interphase library, 9,336 entries in the metaphase library, and 8,139 entries in the MB library. These groups overlapped, giving 7,986 entries with at least 100 reads in all three libraries.

An enrichment threshold of ≥2 was used to identify MB-specific and MB-enriched transcripts. MB-specific transcripts had an RPKM score of ≥2 for both the MB/metaphase and MB/interphase ratios. MB-enriched transcripts had a score of ≥2 in either the MB/metaphase or MB/interphase ratios. The log_2_ enrichment score of MB/metaphase transcripts was plotted against the log_2_ enrichment score of MB/interphase transcripts using the R programming language and package ggplot2 (R Core Team, 2018; https://www.R-project.org/) (*102*)

### Gene ontology

Gene ontology analysis was performed using human ortholog UniProt IDs as input for PANTHER(*103*). Biological process terms (transcription, cell cycle, RNA processing, cell fate, signal transduction, and DNA processing) were assigned through a combination of PANTHER/UniProt analysis and manual annotation and were assembled into Fig. 1 and Supp. Tables 1, 2, and 3.

### Tableau visualization

We delivered our annotated data for the 22 MB-enriched RNAs into Tableau (https://www.tableau.com) to create Fig. 1C. Each color represents an association with a particular gene ontology term, and the size of each circle correlates to the enrichment score.

### Immunofluorescence

MBs and MBRs from synchronized and asynchronized cells (HeLa or CHO), respectively, were fixed. Cells were cultured on high-precision cover glasses, fixed in 3% paraformaldehyde (Electron Microscopy Sciences, Cat# 15735-85) with 0.3% Triton^®^ X-100 (Sigma-Aldrich, Cat# T9284) in PHEM buffer (60 mM PIPES, 27 mM HEPES, 10 mM EGTA, 4 mM MgSO4, pH 7.0) for 10 min at room temperature, blocked for 60 min in blocking solution (PHEM with 3% bovine serum albumin(BSA) (Sigma-Aldrich, Cat# A2153)), and incubated with primary or secondary antibodies in blocking solution (PHEM with 3% BSA). Cover glasses were mounted on slides using Fluoro-Gel mounting medium (Electron Microscopy Sciences, Cat# 17985-03) for SIM microscopy.

### RNAscope/Fluorescent in situ hybridization

RNA in situ hybridization was performed using the RNAscope Multiplex Fluorescent kit (Cat# 323100; Advanced Cell Diagnostics, Inc). in accordance with the manufacturer’s instructions. Briefly, CHO or HeLa cells were fixed for 30 minutes with 4% paraformaldehyde in 0.1 M PBS (15735-85; Electron Microscopy Sciences) on cover glasses coated in poly-L-lysine (P4832; Sigma), dehydrated through a graded ethanol series (50%, 70%, 100%, 100%), and stored overnight at 4°C. Cells were rehydrated through a graded ethanol series (100%, 70%, 50%, PBS, PBS) at room temperature and pretreated with hydrogen peroxide and then protease III for 10 minutes each prior to hybridization. Cells were hybridized using custom RNAscope probe sets designed against *Klf4, Jun, PolyA, Zfp36, Kif23*, and DapB (control) mRNA sequences (Cat# 563611; 563621; 318631; 563631; 558051; 310043; Advanced Cell Diagnostics, Inc., respectively).The preamplifier, amplifier and HRP-labeled probes were then hybridized sequentially, followed by immunofluorescence labeling with Alexa488 or Alexa568 conjugated tyramide (AAT Bioquest, Cat# 11070; Thermo Fisher Scientific Inc, Cat# B40956, respectively). Subsequent immunofluorescent stainings were performed using anti-α-tubulin and/or anti-MKLP1 antibodies in Dulbecco’s PBS (DPBS) with 3% bovine serum albumin and 0.1% saponin (AAA1882014; Thermo Fisher Scientific). Cover glasses were mounted on microscope slides using Fluoro Gel mounting medium.

### Structured illumination microscopy imaging

Structured illumination microscopy was performed on a motorized inverted Eclipse Ti-E structured illumination microscope (Nikon) at the University of Wisconsin–Madison Biochemistry Optical Core. Images were captured on an Andor iXon 897 electron-multiplying charge-coupled device camera (Andor Technology). Images were captured and processed using NIS-Elements AR with N-SIM software (Nikon).

### Hexanediol treatments

HeLa cells were cultured and synchronized as described above. HeLa cells were released from the S phase block by transfer to a normal medium for 8.25 to 8.5 hours, and contractile ring-mediated early MB stages were visually confirmed. Cells were treated with medium supplemented with 7.5% 1,6-hexanediol (240117; Sigma-Aldrich) for 90 seconds, washed with PBS, and incubated in normal medium. Cells were fixed and processed for immunofluorescence as described above. GFP-MKLP1 expressing HeLa cells were treated with hexanediol as described above, and then processed for time-lapse imaging.

### FRAP experiment

GFP-MKLP1 expressing cells were imaged using an inverted NickonEclipse TiE microscope equipped with a CSU-X1 spinning disk confocal scanning unit (Yokogawa) and a EMCCD Camera (Evolve 512 Delta, Photometrics). Bleaching was performed by scanning 3 iterations of 488 nm excitation throughout the bleaching ROI. Images were acquired every 20 seconds with a x100 1.4 NA PL-APO VC objective lens and MetaMorph software (MDS).

### Puromycin labeling to visualize translation in midbodies

HeLa and CHO cells were cultured and synchronized as described above. MB-stage cells or asynchronous cell populations were treated with medium supplemented with 91 μM puromycin for 4 minutes, washed twice in DPBS, and immediately fixed in 3% paraformaldehyde with 0.3% Triton^®^ X-100 in PHEM buffer on poly-L-lysine-coated cover glasses for 10 minutes. Translation was visualized using anti-puromycin primary antibodies (Millipore Sigma, Cat# MABE343), co-incubated with anti-MKLP1 antibodies (Novus Biologicals, Cat# NBP2-56923) as a marker for midbodies and midbody remnants, and processed as described above. We quantified then the number of puro rings we observed at different stages and plotted this using Excel and GraphPad Prism.

### HPG-ClickIT and OPP-ClickIT experiments

For analysis of newly synthetized proteins, HeLa cells were washed and grown in methionine-free RPMI media (Thermo Fisher Scientific, Cat# A1451701) for 2 hours containing HPG (400 μM; manufacturer’s guideline is 50μM) with/without 9.4μM anisomycin or 335μM cycleheximide (A9789; C1988; Sigma, respectively). After incubation, cells were fixed with 4% paraformaldehyde for 15min, washed with DPBS containing 3% BSA and then 0.25% Triton-X-100 was incubated to the cells for 5min. For the detection of Click-IT HPG, Click-IT reaction cocktail containing the Alexa Fluor^®^ 488 azide or BP Fluor 555 azide (BroadPharm, Cat# BP-25564) was incubated for 30min in dark. Additionally Click-IT^®^ Plus OPP Alexa Fluor^®^ 488 protein synthesis assay kit was used for another type of detection of newly synthetized proteins. HeLa cells were incubated in growth media with 20μM Click-IT^®^ OPP (O-propargyl-puromycin) working solution for 4 min, and then fixed by the same methods with HPG Click-IT^®^. The fixed cells were then incubated with Click-IT^®^ OPP reaction cocktail for 30 min at room temperature in dark.

### siRNA experiments and genes

HeLa cells were seeded in 6-well plates and cultured in DMEM/high-glucose/GlutaMAX at 37°C under a 5% CO2 atmosphere before transfection. After one-day growth, siRNA transfection (at a final concentration of 80pmols) was performed using lipofectamine^™^ RNAiMAX. 8μl siRNAs and 6μl siRNA transfection reagent were diluted in each 100μl Opti-MEM^™^ media (Thermo Fisher, Cat# 31985062), then mixed and incubated for 5 min at room temperature. Subsequently, the mixtures totally 214μl were added to each well containing cells and 800μl growth medium. The mixture was then incubated in HeLa cells for 15 hours. Following that, new media were replaced to reduce toxicity of transfected reagent and then the transfected cells were cultured up to 24h or 48h since the transfection. For a matured midbody synchronization, the siRNA transfected cells were blocked by nocodazole (25ng/ml) for 5 hours, then mitotically rounded cells were physically shake with new culture media and the floating cells were transferred on poly-L-lysine coated cover slip. And then the prophase cells were released to the midbody-stage for 4 hours. The synchronization was performed before 9 hours from a fixed time point (24h or 48h).

### Quantification of siRNA experiments

To determine the number of bi-nucleates or multi-nucleates, we visualized DAPI, MKLP1, Phalloidin, and α-tubulin in the control and siRNA treated samples both using 20x objective of an ECHO Revolve Microscope (Echo Laboratories, San Diego, CA, USA). For IST1 siRNA experiments, we determined the number of cells stuck at the midbody bridge stage versus non-dividing cells. All visualized images were analyzed from at least 100 nuclei per each group.

### Quantification of fixed midbody bridges and midbody remnants

To quantify MKLP1, RBP, midbody factors, and tubulin signals in the intercellular bridges or MBRs, we performed line scans using profile plot analysis in ImageJ/FIJI. Fluorescence intensity values represent the average fluorescence intensity measured from a 6.5μm wide line along the axis of the midbody bridge. Statistical analyses were performed using Excel and Graphpad Prism software. All line scan results are shown as mean ± SEM of at least five images.

### Quantification of translation signals in the midbody

To quantify the α-Puro, OPP Click-IT^®^ and HPG(L-Homopropargylglycine) Click-IT^®^ signals we performed line scans. Fluorescence intensity values represent the average fluorescence intensity measured from a 6.5μm wide line along the axis of the midbody bridge. Graphs were assembled using GraphPad Prism. All line scan results are shown as mean ± SEM of at least five images.

### Quantification of FRAP images

To quantify MKLP1 and MKLP2 dynamics in HeLa cells, the mean intensity values for two different regions of identical areas were obtained for each image frame, as photobleached (*F_p_*) and not photobleached (*F_o_*). An empty region of the frame was used to measure the background (*F_b_*). The pre-photobleaching value was normalized to 1 for each sample. The fraction of fluorescent recovery for each frame was calculated as follows: (*F_p_*–*F_b_*)/(*F_o_*–*F_b_*) and plotted as a function of time.

---

**Supplemental Figure S1:**
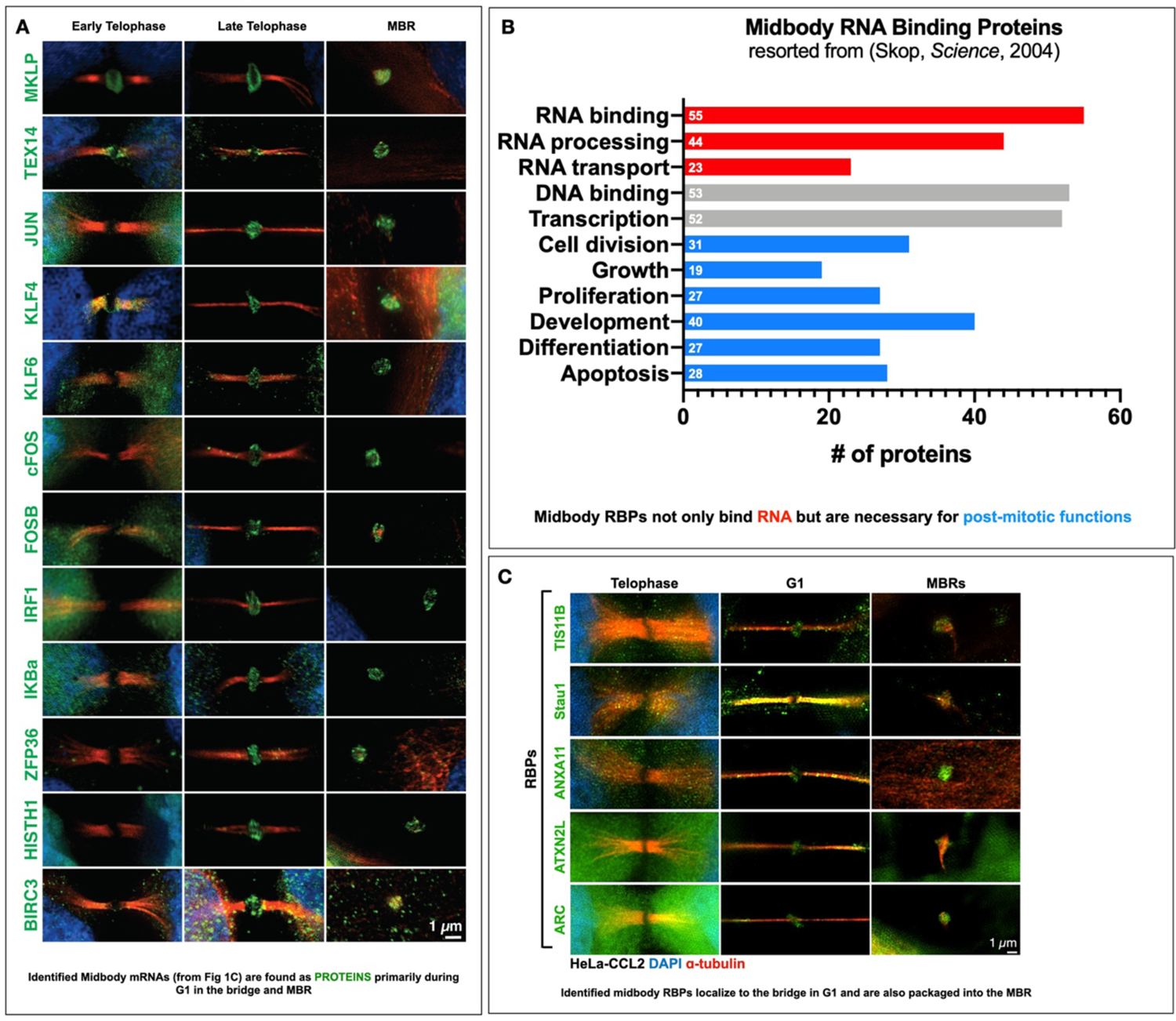
The midbody proteome reveals 99 RNA-binding proteins and 22 factors that are specifically found during late telophase/G1 and in MBRs. (A) Previously identified Midbody RBPs localize to MB during G1 and are packaged into the MBR. We observed puncta along the microtubules as well as in the MB dark zone. Stau, however, is found primarily along the bridge, in a faint ring where we also see ribosomes, and is found in the MBR. (B) MB proteome analyses re-sorted just for RNA binding proteins from in the Supplemental data tables in Skop et al., 2004, identified 99 RNA-binding proteins. When binned by gene ontology terms, many functions in addition to RNA-binding were identified, including DNA binding, cell division, growth, proliferation, development, differentiation, and apoptosis. (C) Of the 22 MB-enriched mRNAs in Fig. 1C, most (n=10/12 assayed) encode proteins that were undetectable until the G1 transition. These include many transcription factors with no known role in cytokinesis, in an environment lacking DNA, suggesting a role in post-mitotic function. MKLP1 and TEX14 have known roles in cytokinesis and were present throughout the MB stages.

**Supplemental Figure S2:**
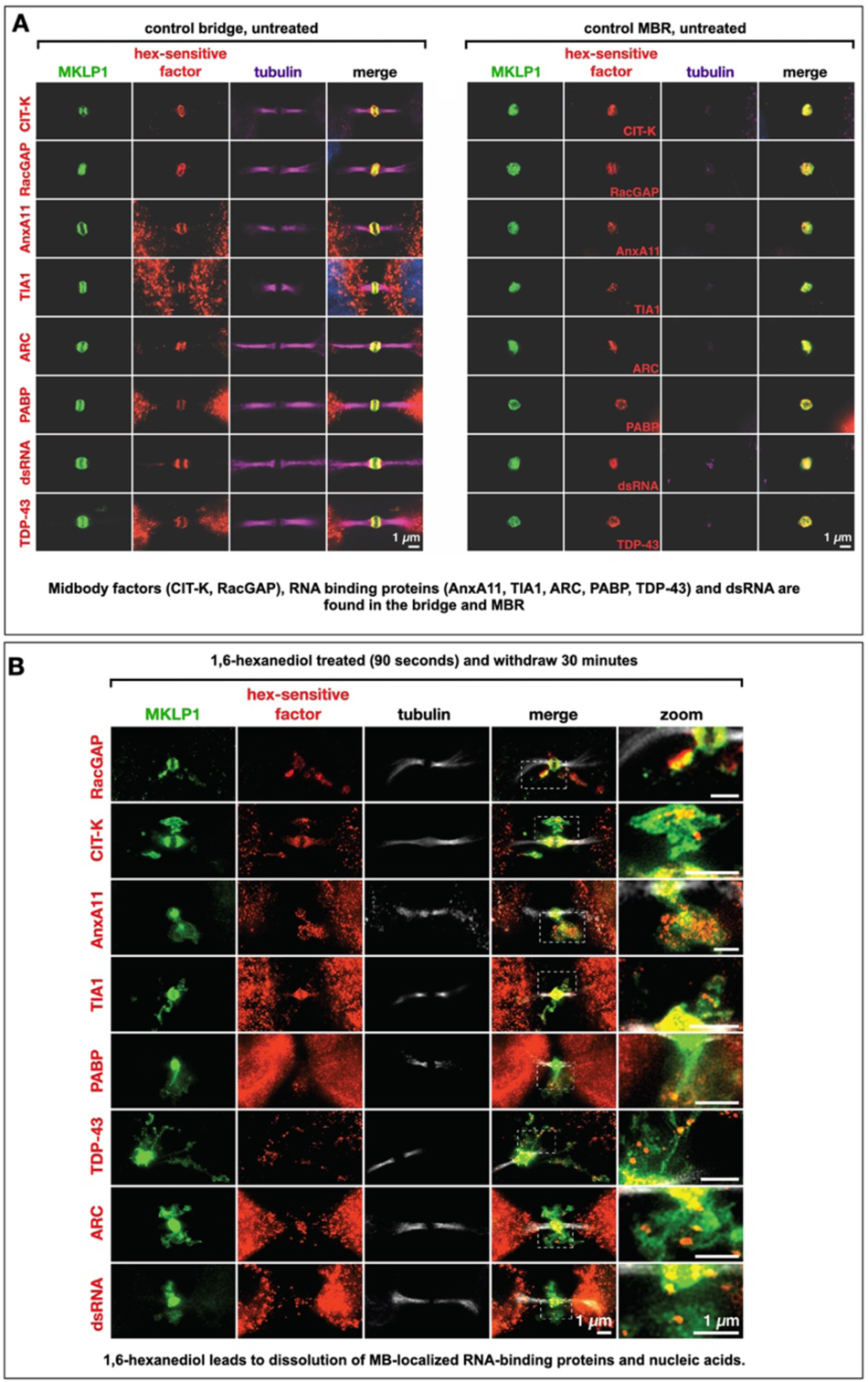
Hexanediol-sensitive proteins and double-stranded RNAs localize to the midbody matrix and are sensitive to hexanediol. **(A)** A range of MB-localized proteins and double-stranded RNAs exhibited sensitivity to 1-6’ hexanediol (hex) treatment, causing their dispersal and progressive reaggregation over time. These factors localized to the MB matrix (red) in mitotic MBs. MKLP1 (green) was used as a marker of the MB matrix, and alpha-tubulin staining (magenta) was used to visualize the dark zone interruption. **(B)** The factors in (A) all remained co-localized with MKLP1 following abscission and release of the MB as an MBR. Scale bars: 1 μm.

**Supplemental Figure S3:**
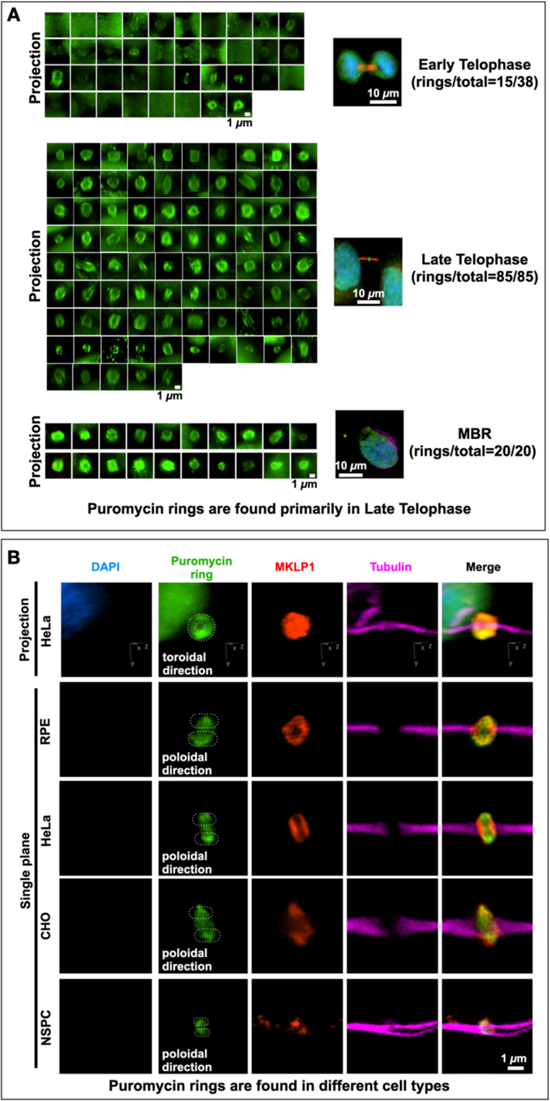
Quantification of the number of puromycin-positive rings during the late stages of mitosis and in different cell types. **(A)** Quantification of the number of distinct puromycin rings observed at different points during the late stages of mitosis, namely early telophase (ET), late telophase (LT), and MBR. The α-Puro label was primarily found in late telophase/G1 and continued in the MBR stage after MBR release. **(B)** Retinal pigment epithelium cells (RPE), HeLa cells, CHO cells, and neural stem/progenitor cells (NSPCs) all had puromycin rings labeled with MKLP1 within the bridge (tubulin). DAPI: 4’,6-diamidino-2-phenylindole. Scale bar: 1 μm

**Supplemental Figure S4:**
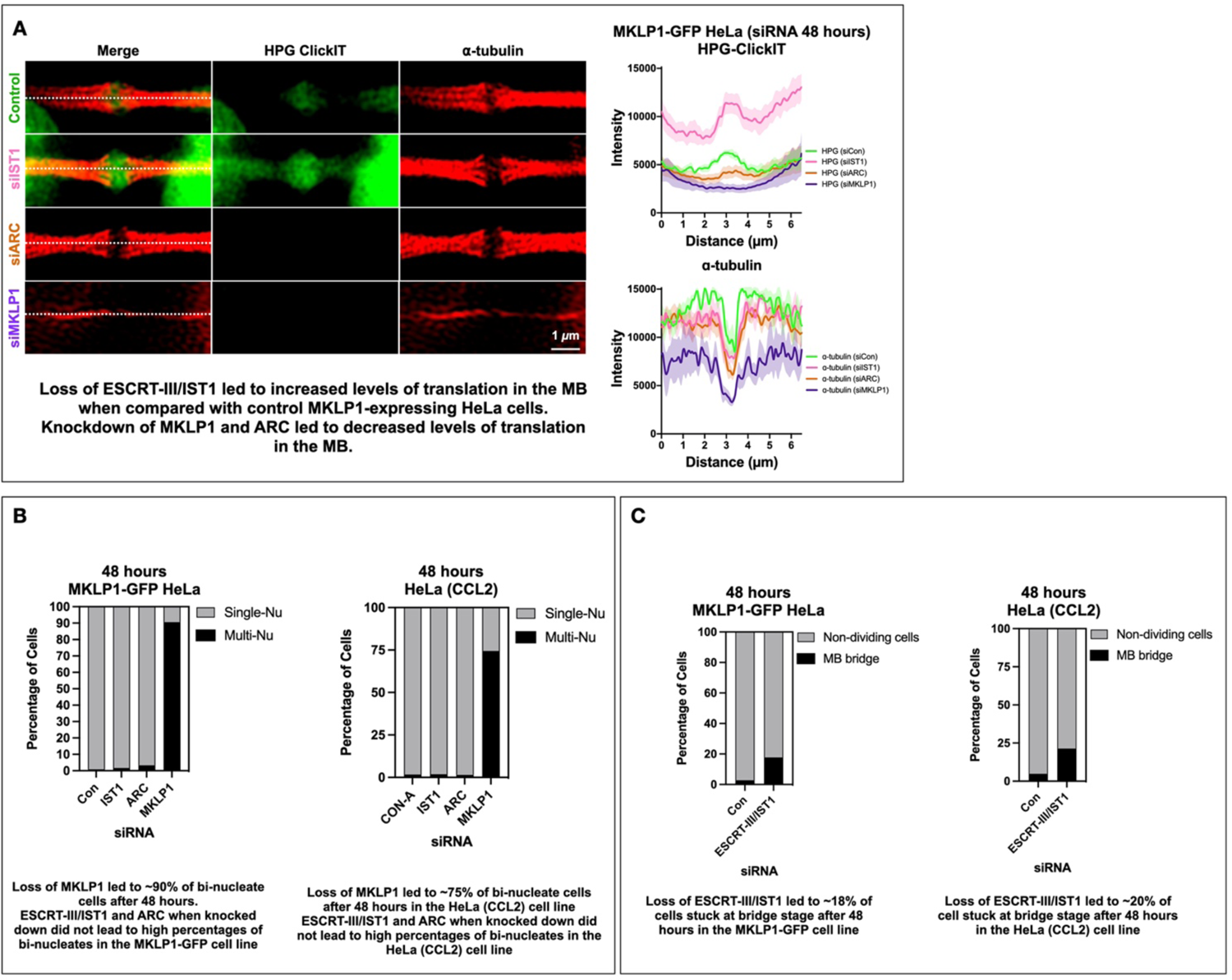
Comparison of siRNA knockdown in HeLa (CCL2) cells and in HeLa cells overexpressing MKLP1-GFP. **(A)** Loss of ESCRT-III/IST1 led to increased levels of translation in the MB dark zone and decreased levels of translation when *MKLP1* and *ARC* were knocked down in the MKLP1-GFP-overexpressing HeLa cell line. **(B)** Loss of MKLP1 after *MKLP1* siRNA treatment led to 90% binucleate cells in the MKLP1-GFP-overexpressing HeLa cell line and 75% binucleate cells in the HeLa (CCL2) cell line after 48 hours. *ESCRT-III/IST1* and *ARC* siRNA did not lead to high levels of binucleate cells in either the MKLP1-GFP-overexpressing HeLa cell line or HeLa (CCL2) cells. **(C)** Loss of ESCRT-III/IST1 caused approximately 18% of cells to be stuck at bridge stage after 48 hours in the MKLP1-GFP-overexpressing HeLa cell line and approximately 20% in the HeLa (CCL2) cell line. Con: control.

**Supplemental Figure S5:**
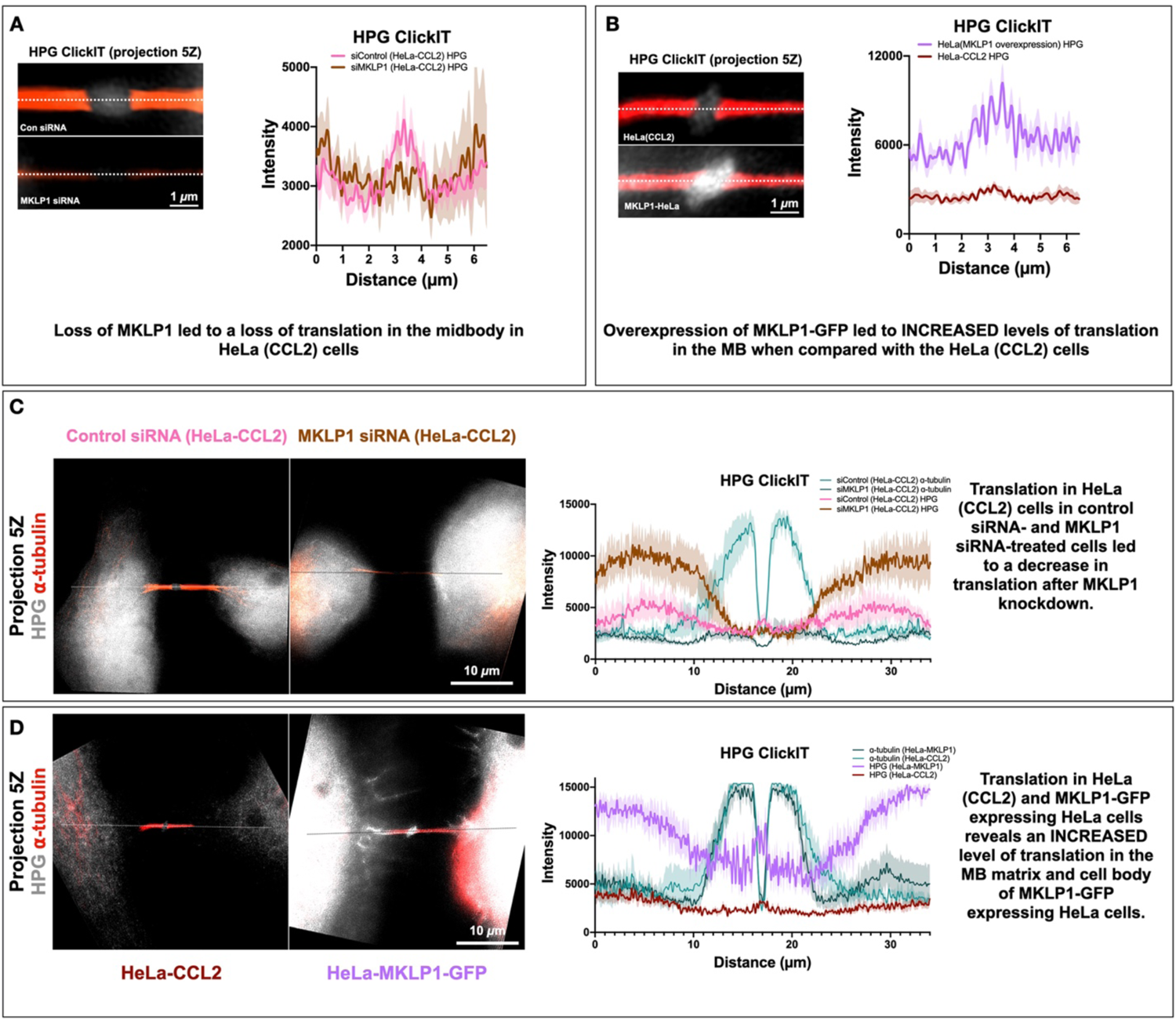
Translation levels are regulated by MKLP1. **(A)** Quantification of the HPG-ClickIT signal in the dark zone of the MB using line scans of the intercellular bridges. A decrease in translation was observed in HeLa (CCL2) native cell lines after MKLP1 knockdown by siRNA treatment, suggesting that MKLP1 may regulate translation in some way. Con: control. **(B)** A remarkable increase in translation was observed in MKLP1-GFP-overexpressing HeLa cells compared with HeLa (CCL2) cells, as visualized by HPG-ClickIT signals. This suggests that MKLP1 itself might regulate levels of translation in general. **(C)** HPG-ClickIT signals were measured in both HeLa (CCL2) cell lines. Translation was localized to the cell body and the MB dark zone. The tubulin scan was removed from the images to show only the translation signal. The tubulin scan is represented in the chart to show where the MB dark zone is located. After MKLP1 siRNA treatment, the signal was lost specifically in the dark zone and not in the cell body. **(D)** To quantify the translation levels in whole cells, HeLa (CCL2) cells and MKLP1-GFP-overexpressing HeLa cells were compared. There was a significant increase in HPG-ClickIT signals in both the cell body and MB in the MKLP1-GFP-overexpressing HeLa cells, suggesting that MKLP1 likely plays a role in mediation of translation levels in general.

**Supplemental Figure S6:**
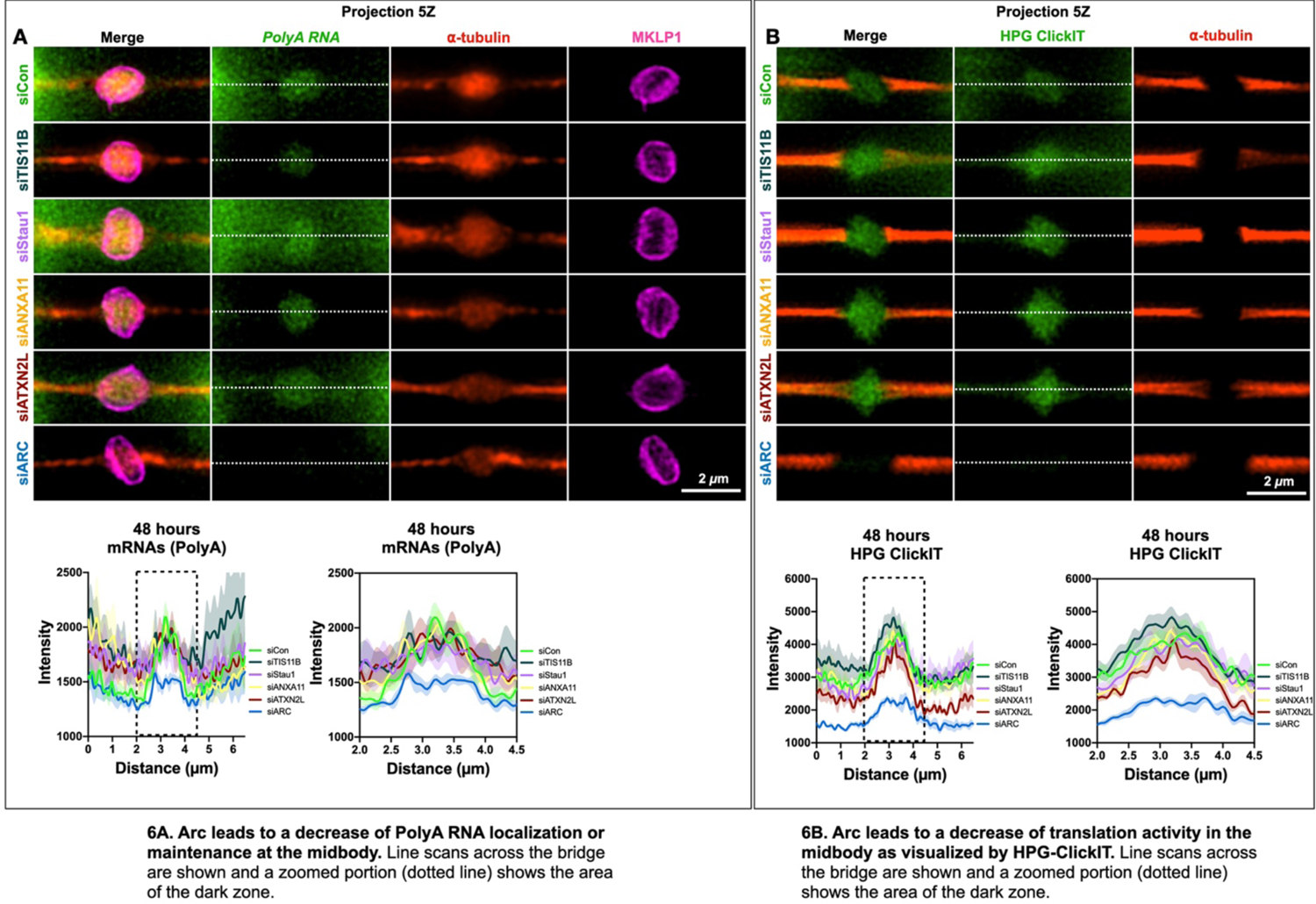
Supplemental Figure S6: Arc is important for PolyA localization or translation at the MB. **(A)** Arc leads to a decrease of PolyA RNA localization or maintenance at the MB. Line scans across the bridge are shown and a zoomed portion (dotted line) shows the area of the dark zone. **(B)** Arc leads to a decrease of translation activity in the midbody as visualized by HPG-ClickIT. Line scans across the bridge are shown and a zoomed portion (dotted line) shows the area of the dark zone.

**Supplemental Table 1: Midbody-specific transcripts.**

MB-specific transcripts were identified as the 22 transcripts that had an enrichment score (RPKM/RPKM) of ≥2 when compared with both interphase (“midbody/interphase enrichment score,” highlighted in yellow) and metaphase (“midbody/metaphase enrichment score,” highlighted in peach). Colors and gene ontology terms correspond to those in Fig. 1C and 1D.

**Supplemental Table 2: Midbody-enriched transcripts in metaphase.**

MB-enriched transcripts in metaphase were identified as the 86 transcripts that had an enrichment score (RPKM/RPKM) of ≥2 when compared with metaphase (“midbody/metaphase enrichment score,” highlighted in peach). Colors and gene ontology terms correspond to those in Fig. 1C and 1D.

**Supplemental Table 3: Midbody-enriched transcripts in interphase.**

MB-enriched transcripts in interphase were identified as the 1051 transcripts that had an enrichment score (RPKM/RPKM) of ≥2 when compared with interphase (“midbody/interphase enrichment score,” highlighted in yellow). Colors and gene ontology terms correspond to those in Fig. 1C and 1D.

**Supplemental Table 4: Raw data from CHO cell RNA-Seq.**

Raw data from RNA-Seq experiments on all three CHO cell libraries (interphase, metaphase, and MB). The CHO RefSeq accession number, the read count, and the RPKM for each transcript in each library are displayed. The human UniProt ID corresponding to each transcript is displayed together with the e-value for each ID. MB/interphase and MB/metaphase enrichment scores and the enrichment scores after log_2_ transformation are given. *Ints2* (UniProt ID: Q9H0H0) and *Cse1l* (UniProt ID: P55060) were used as controls in the qPCR experiments and are highlighted in yellow at the top.

